# Detection of malaria in insectary-reared *Anopheles gambiae* using near-infrared spectroscopy

**DOI:** 10.1101/533802

**Authors:** Marta F. Maia, Melissa Kapulu, Michelle Muthui, Martin G. Wagah, Heather M. Ferguson, Floyd E. Dowell, Francesco Baldini, Lisa-Ranford Cartwright

## Abstract

Large-scale surveillance of mosquito populations is crucial to assess the intensity of vector-borne disease transmission and the impact of control interventions. However, there is a lack of accurate, cost-effective and high-throughput tools for mass-screening of vectors. This study demonstrates proof-of-concept that near-infrared spectroscopy (NIRS) is capable of rapidly identifying laboratory strains of human malaria infection in African mosquito vectors. By using partial least square regression models based on malaria-infected and uninfected *Anopheles gambiae* mosquitoes, we showed that NIRS can detect oocyst- and sporozoite-stage *Plasmodium falciparum* infections with 88% and 95% accuracy, respectively. Accurate, low-cost, reagent-free screening of mosquito populations enabled by NIRS could revolutionize surveillance and elimination strategies for the most important human malaria parasite in its primary African vector species. Further research is needed to evaluate how the method performs in the field following adjustments in the training datasets to include data from wild-caught infected and uninfected mosquitoes.

## Introduction

Malaria is holding back development in endemic countries and remains one of the leading causes of death in children under 5 years-old in sub-Saharan Africa [1–3]. During the past decade, the large-scale roll-out of long-lasting insecticide treated nets and indoor residual spraying across Africa has resulted in a substantial reduction in malaria cases [4]. The WHO’s Global Technical Strategy for Malaria 2016-2030 seeks to reduce malaria incidence and related mortality by at least 90% and to eliminate the disease in a minimum of 35 countries [1]. These bold goals will require new interventions that can address residual malaria transmission as well as new tools to better monitor their impact on vector-borne disease transmission. Mosquito surveillance is a cornerstone of the control of malaria and other vector-borne diseases [5]. However, presently, there is no high-throughput, cost-efficient method to identify *Plasmodium* infection and infectiousness in mosquitoes. Molecular methods such as ELISA and PCR are used to determine parasite infection, but these are expensive and laborious [6–8], challenging resource-poor countries with few funds and limited access to reagents and equipment, and thus are unsuitable for large-scale surveillance. A further complication is that typically only 1-2% of mosquitoes may be infected with transmission stage parasites (sporozoites), meaning that very large sample sizes must be tested to accurately quantify site and time-specific estimates of mosquito infection rates as will be required to assess progress towards malaria elimination [9].

Recent advances indicate several mosquito traits can be accurately identified through analysis of their tissues with near infrared spectroscopy (NIRS) [10–13]. Here, visible and NIR light (wavelength 400-2500 nanometers) is passed through the whole or part of a mosquito specimen and an absorbance spectrum is collected instantly without destroying the sample. Changes in spectral peaks at different wavelengths represent how intensely different molecules absorb light, and thus NIR spectra of mosquitoes are determined by the biochemical composition of their tissues, which are known to differ according to age [14, 15], species [16, 17], microbiome [18], physiological stage [19, 20], and by pathogen infection [20, 21]. Differences in NIR spectra have been used to distinguish young (e.g. <7 days old) from older (7+ days old) malaria vectors, to identify morphologically identical *Anopheles* sibling species, and to detect the presence of the endosymbiont *Wolbachia* in *Aedes aegypti* mosquitoes [10–12]. Most recently, NIRS has been used to detect rodent malaria infections in laboratory-reared *Anopheles stephensi* mosquitoes [22] and Zika virus in *Ae. aegypti* mosquitoes [23]. The use of NIRS has not previously been investigated on human malaria infected mosquitoes. The presence of the parasite-specific proteins and other biochemical changes induced by malaria infection in the vector may permit these to be distinguished from uninfected mosquitoes using spectral tools such as NIRS [24, 25]

Parasite infection in the mosquito can be found in two main forms defined by their parasite development stages: midgut oocyst infections occurring around 2-8 days after feeding on infectious blood; and sporozoite infections occurring 9-14 days after infection, characterized by the release of sporozoites from oocysts into the mosquito’s haemocoel and salivary glands, enabling the mosquito to infect the next human host. Given the different nature of the two infection stages the NIRS profile of an oocyst-infected mosquito may not be the same as a sporozoite-infected one. For this reason, we aimed to test whether NIRS could successfully identify oocyst and sporozoite infections in *Anopheles* vectors, and estimate if the method’s prediction accuracy is dependent on the intensity of infection in the mosquito.

In this paper, we present the successful application of NIRS to differentiate *Plasmodium falciparum*-infected mosquitoes from uninfected mosquitoes, providing the first evidence of detection of human malaria infections in the *A. gambiae* mosquito vector by this cost-effective, fast and reagent-free method. The development of a tool such as NIRS to measure malaria infection rates in mosquito populations would be of great service to malaria pre-elimination efforts as it would allow the processing of large numbers of mosquitoes increasing the accuracy of the estimates of human exposure to malaria infection across different regions, and advancing malaria vector surveillance in Africa.

## Results

### Experimental infections

Approximately 750 female *A. gambiae* (Keele line) [26] of different ages (3-6 days old) were offered a blood meal containing NF54 gametocyte cultures in standard membrane-feeding assays (SMFAs) in three independent replicate experiments. Control (uninfected) mosquitoes were generated by feeding approximately 450 mosquitoes the same blood after gametogenesis was completed. Both groups were represented with mosquitoes of similar ages, between 3 and 6 days old (Table 1 and 2 of supplementary information). Mosquitoes were maintained for 7 and 14 days under insectary conditions to allow oocyst (D7) and sporozoite (D14) development, on each day of sampling live mosquitoes were removed, killed and immediately scanned using NIRS.

**Table 1.** Description of the gametocytemia used for each of the three standard membrane feeding assays (SMFA), number of days kept post blood feeding, number of mosquitoes processed by quantitative PCR (qPCR), % prevalence, and the intensity of infection described as the median and interquartile range (IQR) of the number of parasite genomes present in infected mosquitoes, excluding mosquitoes with no infection.

| SMFA | Estimated gametocytemia | Day post-infectious blood meal | No. mosquitoes tested (n=634) | Positive (N=423) | % prevalence of infection | Intensity of infection: median number of parasite genomes and IQR. |
| --- | --- | --- | --- | --- | --- | --- |
| 1 | 1% | 7 | 175 | 105 | 60.0% | 680 (283-1625) |
|  |  | 14 | n.d. | n.d. | n.d. | n.d. |
| 2 | 0.1% | 7 | 104 | 73 | 70.2% | 456 (67-2052) |
|  |  | 14 | 99 | 47 | 47% | 516 (211 – 6081) |
| 3 | 0.1% | 7 | 114 | 85 | 74.6% | 2995 (3210-8881) |
|  |  | 14 | 142 | 113 | 80% | 10114 (2540 – 29145) |

**Table 2.** Generalised linear mixed-effects models investigating the effect of infection presence (infected or uninfected) and infection load (number of parasite genomes/μl of DNA extract quantified using qPCR) on the PLS score of the predicted samples including mosquito age as a random effect.

|  | Coefficient<br>s | Robust<br>standard error | z | 95% Confidence<br>intervals | P value |
| --- | --- | --- | --- | --- | --- |
| **Oocyst infections** |  |  |  |  |  |
| Infection presence | 0.67 | 0.13 | 5.11 | 0.41 to 0.93 | <0.001 |
| Infection load | -0.000003 | -0.000002 | -1.26 | -0.0000074 to 0.0000015 | 0.21 |
| *Mosquito age*<br>(random effect) | .018 | .005 | - | 0.01 to 0.03 | - |
| **Sporozoite infections** |  |  |  |  |  |
| Infection presence | 0.75 | 0.12 | 6.03 | 0.51 to 1.00 | <0.001 |
| Infection load | 0.0000019 | 0.0000003 | 5.82 | 0.0000013 to 0.0000025 | <0.001 |
| *Mosquito age*<br>(random effect) | 0.007 | 0.008 | - | 0.0032 to 0.087 | - |

Mosquitoes fed on infectious blood were analysed by quantitative polymerase chain reaction (qPCR) for intensity of infection. Additionally, 60 mosquitoes from the control groups (30 from feed 2 and 3 respectively) were also analysed by qPCR to confirm the absence of malaria infection. No mosquitoes from these control groups tested positive for infection.

The minimum number of parasite genomes detectable per mosquito was 10 parasite genomes/ per μl of DNA extract, calculated from standard curves generated for each qPCR run using a 5-point 10-fold serial dilution of DNA extracted from asexual NF54 cultures synchronized to ring stage. This gives a threshold detection of ~500 parasite genomes per mosquito for the qPCR assay.

### Near infrared spectra selection

A total of 634 *A. gambiae* (Keele strain) were scanned using NIRS (Table 1). DNA was extracted and analyzed for *P. falciparum* infection by qPCR as described above. Samples with inconclusive qPCR results or poor spectra quality were excluded (n=72). Poor quality or outlier spectra were visually identified by comparing them to all other spectra, and spectra that were prominently flat or prominently noisy were excluded, as described elsewhere [10]. Thus, NIR absorbance spectra and respective infection status data from a final total of 562 mosquitoes were used to estimate the accuracy of NIRS for prediction of malaria infection (Figure 1).

**Figure 1.**
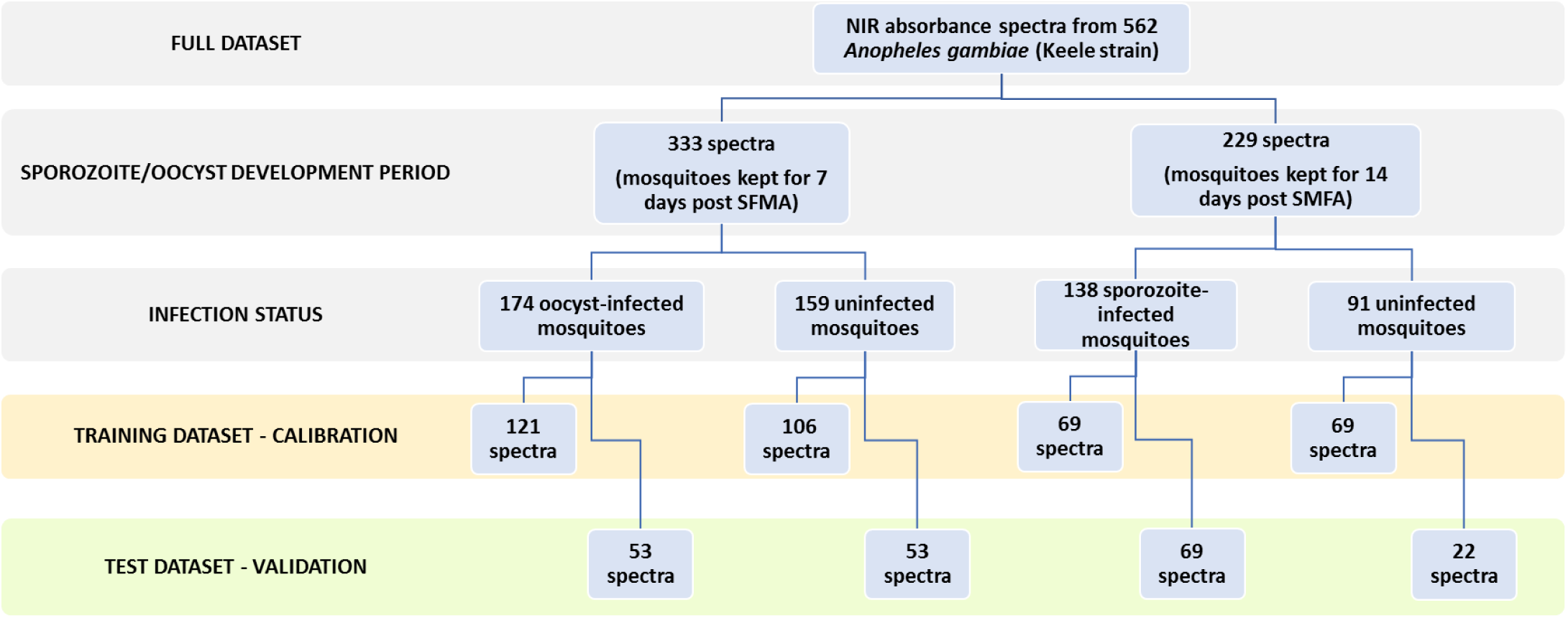
Study flow chart showing number of spectra collected, infection status and random assignment of spectra to either training or test dataset.

### Model prediction accuracy

The relationship between spectra and infection was analyzed using partial least square regression (PLS). Training datasets were used to perform multiple leave-one-out cross validations (LOOCV) and develop two calibrations, one for prediction of oocyst infection and another for prediction of sporozoite infection. The calibrations were then validated using test datasets composed of samples with unknown infection status that had not been included in the calibration’s training dataset. The number of factors used in the calibration was 12, determined from the prediction residual error sum of squares (PRESS) and regression coefficient plots (see supplementary information). In the PLS model, a value of “1” was assigned to all the actual uninfected samples whereas a value of “2” was assigned to the actual infected mosquitoes (infection as defined by the qPCR results). The PLS calibration derived components used to transform the original spectra of each predicted independent sample into a PLS score; a score value of 1.5 was considered as the threshold for correct or incorrect classification, meaning any mosquito with PLS score below 1.5 was predicted as uninfected and equal or greater than 1.5 was predicted as infected. The PLS model showed that NIR spectra from both oocyst and sporozoite infected mosquitoes were distinct from their counterpart uninfected mosquitoes with 91.2% (86.7% − 94.5%) and 92.8% (87.1% − 96.5%) self-prediction accuracy respectively (Figure 2a and 3a). When tested on samples with unknown infections status that had not been included in the training dataset, the calibration maintained high sensitivity and specificity at both detecting oocyst and sporozoite infection, with 87.7% (95%CI: 79.9% −93.3%; Cohen’s kappa=0.75)) and 94.5% (95%CI: 87.6% − 98.2%; Cohen’s kappa=0.86) prediction accuracy respectively (Figures 2b and 3b).

**Figure 2.**
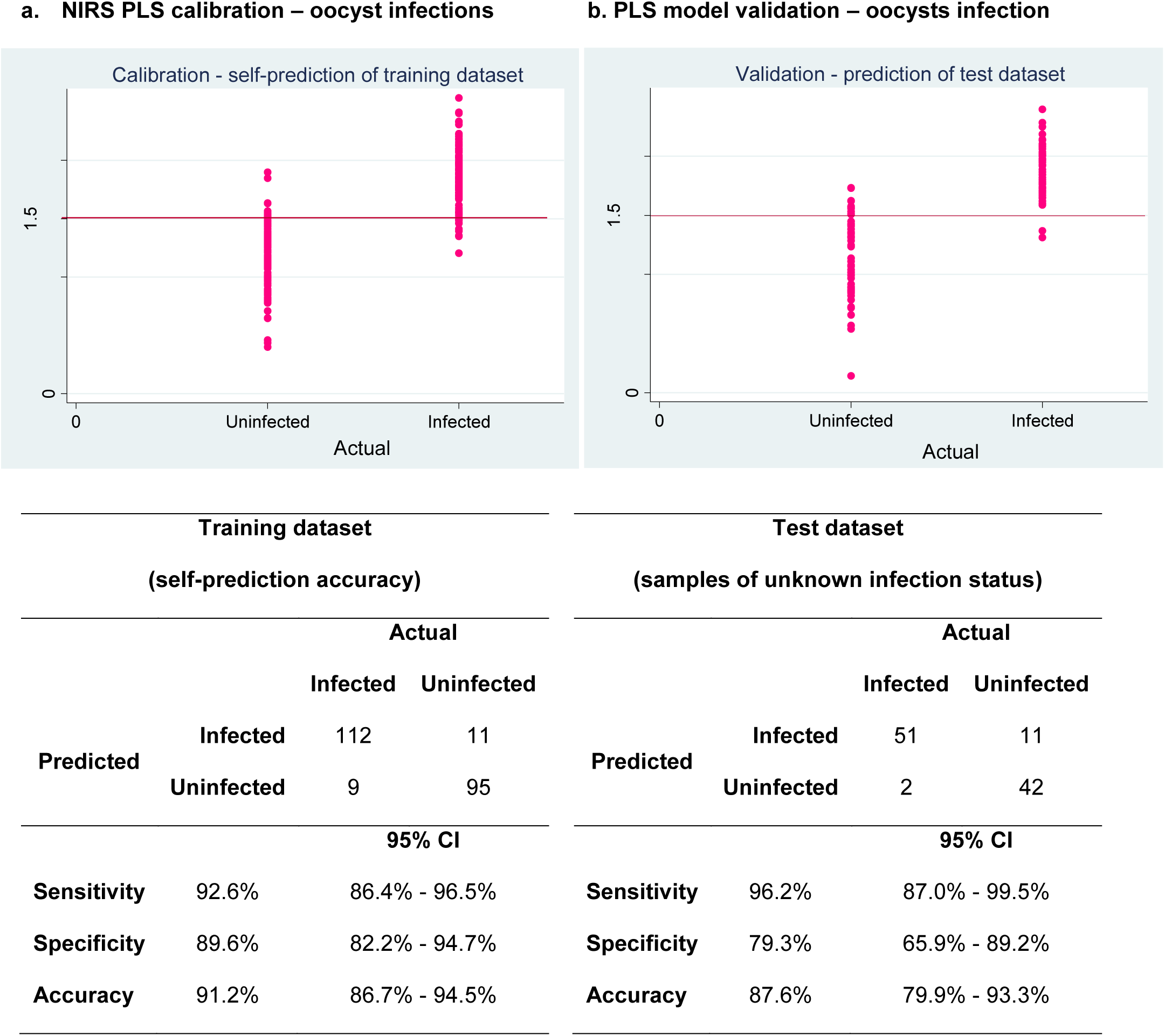
Actual versus predicted plots of oocyst infected mosquitoes investigating NIRS as diagnostic method. Sensitivity, specificity, accuracy and respective 95% confidence intervals of self-prediction of *P. falciparum*-infection in training dataset (left) and prediction of samples of unknown status in test dataset (right) (PLS scores: 1= uninfected, 2= infected and 1.5 as cut-off value).

**Figure 3.**
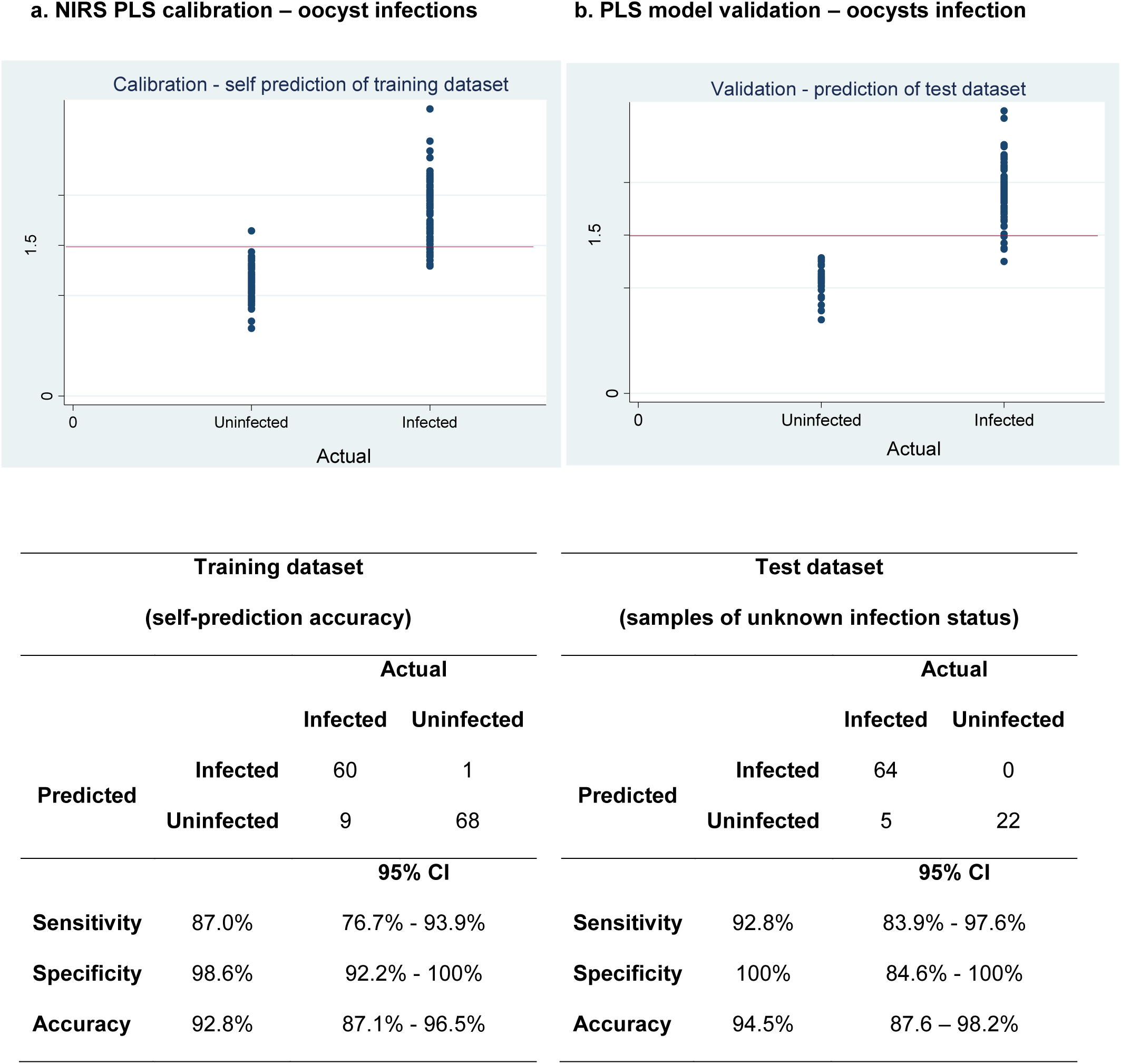
Actual versus predicted plots of sporozoite infected mosquitoes investigating NIRS as diagnostic method. Sensitivity, specificity, accuracy and respective 95% confidence intervals of self-prediction of *P. falciparum*-infection in training dataset (left) and prediction of samples of unknown status in test dataset (right) (PLS scores: 1= uninfected, 2= infected and 1.5 as cut-off value).

### Infection load and prediction accuracy

The parasite load in a mosquito is of epidemiological importance as there is evidence of a continual increase in transmission potential with increasing sporozoites numbers [27]. To test if the NIR prediction output scores were affected by parasite load qPCR was done to estimate the relative number of parasite genomes in each infected mosquito (Table 2) and used to evaluate the calibration model’s accuracy. The oocyst-infected mosquitoes in the test data set had a range of infection loads (Median: 1925, IQR: [295 to 4883]). Two oocyst-infected mosquitoes were misclassified as uninfected, both of which had relatively low infection loads (357 and 389 parasite genomes/μl of DNA extract) (Figure 4a). Generalised linear mixed-effects models were used to investigate the effect of infection load and infection presence on the PLS scores (response variable) of the predicted samples. The age of the mosquitoes on the day of the infectious feed was included as a random effect. It was observed that the presence of oocyst infection influenced the NIRS prediction score (Coefficient: 0.67; 95%CI: 0.41 to 0.93; p<0.001) but the infection load did not (Coefficient: −0.000003; 95%CI: −0.0000074 to 0.0000015; p-value: 0.21). The sporozoite-infected mosquitoes in the test dataset had a range of infection loads (Median: 8841, IQR: [2516 to 20112]). Five sporozoite-infected mosquitoes were misclassified as uninfected: two presented with the lowest infection loads of the test dataset (33 and 38 parasite genomes/μl of DNA extract); the other three had relatively high infection loads (1156, 6660 and 12591 parasite genomes/μl of DNA extract) (Figure 4b). The presence of sporozoite significantly affected the PLS scores of the predicted samples (Coefficient: 0.75; 95%CI: 0.51 to 1.00; p-value: <0.001) as did the infection load (Coefficient: 0.0000019; 95%CI: 0.0000013 to 0.0000025; p-value<0.001).

**Figure 4.**
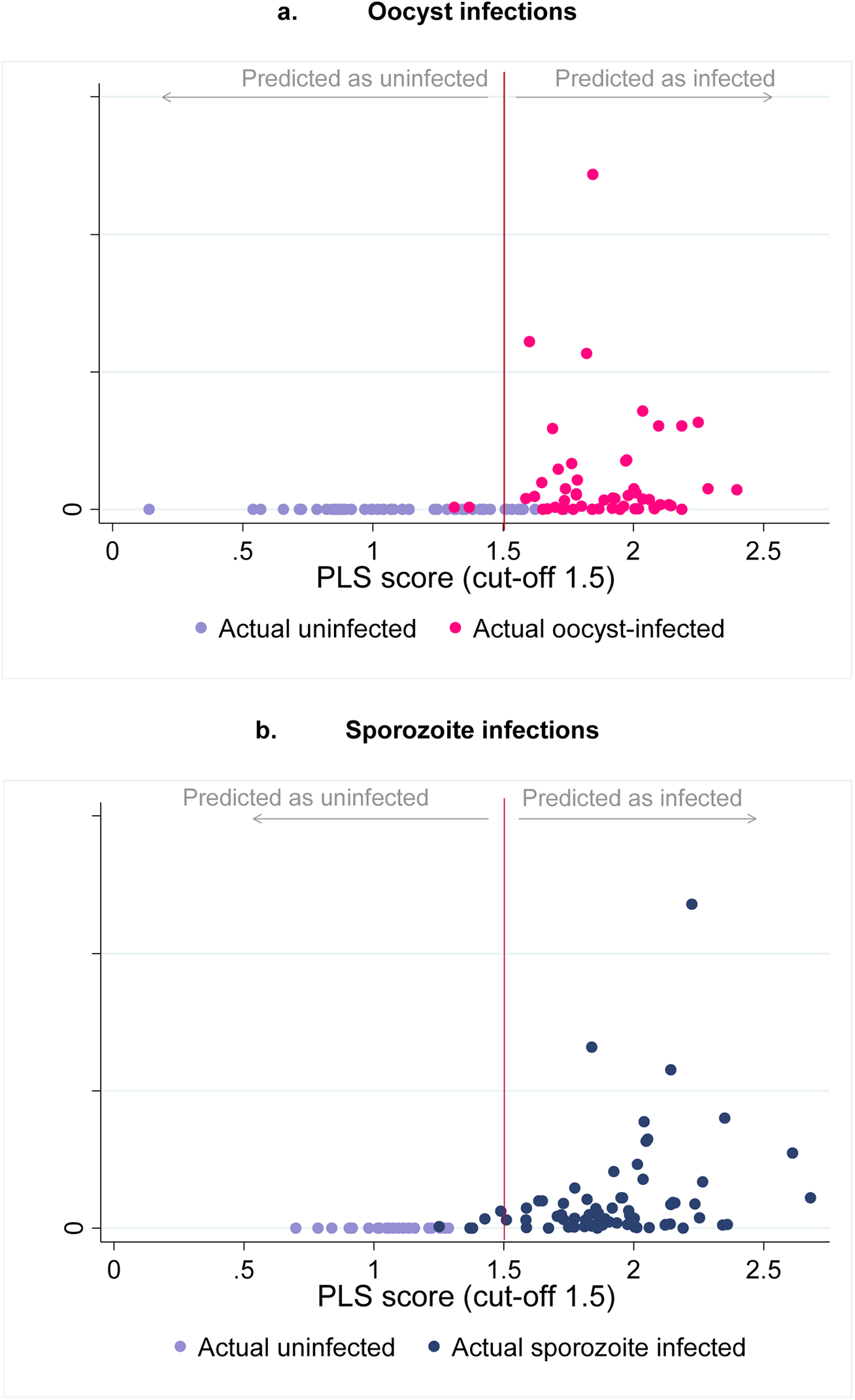
Intensity of *P. falciparum* oocyst (a) and sporozoite (b) infection, quantified as the number of parasite genomes per μl of DNA extract, in *A. gambiae* mosquitoes and prediction value score based on the predicted probability of infection, with 1= predicted as not infected and 2=predicted as infected (cut-off value of 1.5)

## DISCUSSION

This is the first study to show that NIRS can be used to accurately detect human malaria in *A. gambiae* mosquitoes. NIRS predicted oocyst infection with 87.7% accuracy (79.9% − 93.3%) and sporozoite infection with 94.5% accuracy (87.6% – 98.2%). The NIRS predictive accuracy for sporozoite infection of >90% in this study concurs with previous work done using the rodent malaria in *Anopheles stephensi*, which found that NIRS could detect the presence of sporozoites in infected mosquitoes with 77% accuracy [22]. Unlike the previous study, the present calibration model was also capable of identifying oocyst-infected mosquitoes. The PLS calibration of the present study was based on a narrower interval of the electromagnetic spectrum, 500 to 2400 nm, compared to 350 to 2500 nm. This narrower range excludes noise present in the extremities of the spectra due to light source and sensor limitations and therewith improved the prediction accuracy of the calibration model. Furthermore, the previous study used spectra from mosquitoes that had been saturated with chloroform which was used to knock them down. This contamination led to clear chloroform peaks in the NIR spectra which may have added to the noise and reduced prediction accuracy of the calibration. Differences between the vector species and parasite species may also have played a role in the small discrepancy of predictive accuracy between studies. In addition, the experimental approach used in the present study, also permitted to account for the potentially confounding effects of the infected bloodmeal, given that control group had been fed the same blood but with inactivated gametocytes.

Near infrared light is absorbed differently by diverse biochemical compounds which, in the mosquito, may consistently vary with between species, age and in this case infection status. It is hypothesized that biochemical changes occurring in the mosquito, as a consequence of *P. falciparum* infection, made it possible to distinguish between infected and uninfected mosquitoes using NIRS. Consistent differences between the NIR absorbance spectra of infected and uninfected mosquitoes may be related to the presence of parasite-specific molecules in the infected mosquitoes [28–30]. Also, it is possible that tissue changes may occur in the mosquitos due to their immune response to the parasite which could have an effect on the biochemical composition of the mosquito [28]. Additionally, it is known that *Plasmodium* infection alters metabolic pathways in mosquitoes and leads to higher energy resource storage [31] which may lead to differences in NIRS spectra. More research is needed to better understand the underlying biochemical features that enable NIRS to distinguish between *Plasmodium*-infected and uninfected mosquitoes.

The prediction accuracy of the NIRS calibration to detect sporozoite infection was influenced not only by the presence of *Plasmodium falciparum* sporozoites but also the parasite load (number of parasite genomes). This was not the case of the calibration to detect oocysts, which was only significantly influenced by the presence of infection in the midgut. It is possible that slight differences in DNA extraction efficiency between samples may have affected the estimate number of parasite genomes in each insect sample and therefore it is imprudent to make conclusions on how strongly infection load may be influencing the PLS output scores. The performance accuracy of NIRS was similar to qPCR (sporozoite detection: Cohens kappa=0.86; oocyst detection: Cohens’s kappa=0.75). The strong inter-rate agreement between the two methods, suggests that NIRS may have similar sensitivity and specificity as qPCR at detecting malaria sporozoites in the mosquito host. ELISA is less specific than PCR [32], however due to its low-cost and ease, it is routinely the assay chosen by surveillance programs to measure the proportion of mosquitoes that carry sporozoites and the entomological inoculation rate (EIR). It is possible that EIR estimates could be improved by using a more accurate diagnostic test. However, a direct comparison of NIRS and ELISA was not the objective of this study. Presently NIRS still requires further optimization and validation in the field before being considered as a possible replacement for ELISA in surveillance programs. While the results presented in this paper are promising, NIRS calibrations generated using lab-reared mosquitoes do not necessarily represent the diversity of vectors in the field, providing no guarantee of the robustness of the method when tested on wild-caught mosquitoes. Calibrations must be based on training datasets that capture the diversity of field-mosquitoes reducing confounders that may affect the classification accuracy, including, different mosquito species, age, infection, size, insecticide resistance status, microbiome, and origin. NIRS is a promising technology that may provide an accurate and high-throughput solution to monitoring malaria transmission in the vector as progression towards elimination is made. Such a tool may revolutionize how entomological data is used by control and research programmes given that the same test can report various entomological parameters, including age, species and infection status, therewith compiling vast information of epidemiological importance to understanding how vector populations and malaria transmission are changing. Future research efforts and resources need to be directed at evaluating the best way of generating and optimizing calibrations based on wild-caught mosquitoes for each entomological parameter, and validating these using specimens from different ecological and geographical regions.

## MATERIALS AND METHODS

### Mosquitoes

Mosquitoes from a colony of *A. gambiae* (Keele line) [26] were reared under standard insectary conditions (26±1°C, 80% humidity, 12 hr light:12 hr dark cycle) at the University of Glasgow, Scotland, UK. Larvae were fed on Tetramin tropical flakes and Tetra Pond Pellets (Tetra Ltd, UK). Pupae were transferred into cages for adult emergence. Adult mosquitoes were fed *ad libitum* on 5% glucose solution containing 0.05% (w/v) 4-aminobenzoic acid (PABA). SMFA was done with 3-6 days old mosquitoes.

### Parasite culture and standard membrane feeding assays (SMFA)

*P. falciparum* (NF54) parasites were cultured using standard methodology to produce infectious gametocytes [33], using human blood and serum obtained from the Glasgow and West of Scotland Blood Transfusion Service. Standard membrane feeding assays (SMFA) were conducted on three different occasions using gametocytes produced *in vitro*: the first SMFA was done with a high gametocyte density (approx. 1% gametocytes) and the two-subsequent feeds with a lower density (~ 0.1% gametocytes) to produce more uninfected mosquitoes. For each SMFA, 300 female *A. gambiae* s.s (Keele line) mosquitoes 3-6 days post emergence were distributed in pairs into 6 cups of 50 mosquitoes each. In the first SMFA, mosquitoes were 3,4 and 5 days old, in the second SMFA they were 4,5 and 6 days old and in the third SMFA mosquitoes were 3 (2 pairs of cups) and 4 days old. One cup of each pair was offered blood with infectious gametocytes and allowed to feed for 20 minutes. The temperature of the membrane feeders was then reduced to below 30°C for 30 minutes to allow all mature gametocytes to complete gametogenesis [34]. The remaining cups of mosquitoes were then allowed to feed on the same blood, to produce control mosquitoes with zero infection rates, and thus obtain a comparable control sample differing only in the complete absence of parasite infection.

### Near infrared spectra collection and data analysis

After feeding, the blood-fed mosquitoes in each pot were maintained for 14 days under insectary conditions and examined for oocyst and sporozoite development on day 7 and 14 days post infection respectively. Mosquitoes were killed using chloroform vapour before collecting near infrared absorbance spectra from each individual mosquito without any further processing, using a Labspec 4i NIR spectrometer with an internal 18.6 W light source (ASD Inc, Longmont, CO) and ASD software RS^3^ per established protocols [10], but using a 3.2 mm-diameter bifurcated fibre-optic probe which contained a single 600 micron collection fibre surrounded by six 600 micron illumination fibres. The probe was placed 2.4 mm from a spectralon plate onto which the mosquitoes were placed for scanning. Spectra between 500–2400 nm were analysed through leave-one-out cross validations (LOOCV) using partial least square (PLS) regression in GRAMS Plus/IQ software (Thermo Galactic, Salem, NH). After scanning, each mosquito carcass was stored individually at −80 °C in ATL lysis buffer (QIAGEN) until DNA extraction, to perform qPCR to determine the infection status of the mosquito.

### DNA extraction and quantitative real-time polymerase chain reaction (qPCR)

DNA was extracted using Qiagen DNeasy Blood & Tissue^®^ DNA extraction kits from mosquito abdomens (for mosquitoes analyzed 7 days post infectious feed) and whole mosquitoes (for mosquitoes killed 14 days post infectious feed) and eluted in 50 µl of water. A 20 µl aliquot of the 50 µl of extracted DNA for each mosquito was transferred to individual wells of DNAstable^®^ 96 well plates (Sigma-Aldrich) and allowed to air dry at room temperature. The plates were shipped to KEMRI Wellcome Trust (Killifi, Kenya) for qPCR analysis. Samples were reconstituted in 20 µl of DNAse-free water and *P. falciparum* genome numbers present were quantified by qPCR. Quantification reactions were performed in 15 μL volumes, containing 1.2 μl of 10 mM forward and reverse primers (377F: 5’ ACTCCAGAAGAAGAAGAGCAAGC-3’; 377R: 5’-TTCATCAGTAAAAAAAGAATCGTCATC-3’; 7.5 μL of SYBR^®^ Green PCR Master Mix, 1.1 μL of DNAse-free water and 4 μL of sample DNA, using an Applied Biosystems 7500 Real-Time PCR System. The cycling profile comprised an initial denaturation of 95 °C for 900 s (holding stage) and then 40 amplification cycles of denaturation 95°C for 30s (seconds), annealing 55°C for 20s and extension 68°C for 30s. At the end of amplification, melt curves were produced with 15s denaturation at 95°C, followed by 60s at 60°C, 30 s at 95°C and 15 s at 60°C. Parasite load was estimated for each sample by comparison with the standard curve drawn from the DNA standards using Applied Biosystems 7500 software v2.0.6. Samples which amplified after 38 cycles, or which showed a shift in melt curve or two melt curve peaks were excluded.

DNA extracted from uninfected mosquitoes (abdomens and cephalothorax) were used as negative controls, in addition to negative controls with no DNA. Standard curves were generated for each qPCR run using a 5-point 10-fold serial dilution of DNA extracted from asexual NF54 cultures synchronized to ring stage, starting with 100,000 parasites/μl (100,000 parasites; 10,000 parasites; 1,000 parasites; 100 parasites and 10 parasites), run in duplicate.

### Analysis using PLS leave-one-out cross-validations (LOOCV)

The *P. falciparum* detection model was trained and tested according to previously published methods[10] using partial least square(PLS) regression to develop a calibration based on a training data set, which was then used to predict the infection status of samples contained in a test dataset and therewith validate the prediction accuracy of the calibration.

Leave-one-out cross validation (LOOCV) was used to determine if NIR spectra of uninfected mosquitoes were distinct from *P. falciparum*-infected mosquitoes, and to give information on the prediction accuracy of the model to distinguish between infected and uninfected mosquitoes. LOOCCV is a *k*-fold cross validation, with *k* equal to *n*, the number of spectra in a training dataset. That means that *n* separate times, the function approximator is trained on all the spectra except for one spectrum and a prediction is made for that spectrum. Multiple LOOCV based on the training dataset were used to develop a calibration file which was then used to test the predictive ability of the model on a spectra collected from a test dataset (Figure 1).

The results from the qPCR were used to identify which individual mosquitoes, that had been fed an infectious blood meal, had confirmed oocyst and sporozoite infections. This information was then specified to each spectrum and these were randomly assigned to either the training dataset or the test dataset whilst ensuring the same proportion of different mosquito ages was found in the training and test datasets. All uninfected mosquitoes were from the group that had been fed blood without viable gametocytes. A total of 69 sporozoite-infected and 69 uninfected mosquitoes that had been kept for 14 days post SMFA were used to perform multiple LOOCV and generate a calibration file. The same was done using spectra from 121 oocyst-infected mosquitoes and 110 uninfected mosquitoes kept for 7 days post SMFA.

Two separate LOOCV were run to investigate the prediction accuracy of oocyst-infected vs. uninfected, and sporozoite-infected and uninfected mosquitoes respectively. The models were run on Grams IQ software (Thermo Galactic, Salem, NH) and a total of 12 latent factors were selected by visualizing the prediction residual error sum of squares (PRESS) curve, and choosing the minimum number of factors needed to reduce the prediction error of the model without overfitting it. Actual vs Predicted plots were drawn by plotting the actual constituent values (coded as 1= uninfected and 2= infected) on the x axis, and model predicted values on the y axis. Prediction values were generated according to previously published methods [10], values below 1.5 were considered to be predicted as uninfected and values equal to or above 1.5 predicted as infected

The mosquito spectra that had not been included in the training dataset used for developing the calibration were randomly assigned to the test dataset to validate the performance accuracy of the model serving as an independent set of samples. The calibrations generated for detecting *P. falciparum* sporozoite and oocyst infection were validated using 69 sporozoite-infected and 22 uninfected, and, 53 oocyst-infected and 56 uninfected, respectively (Figure 1). This was done by generating a calibration with 12 latent factors based on the training dataset which was then loaded into IQPredict software and used to obtain PLS scores of the independent samples based on the predicted probability of infection, with 1= predicted as uninfected, 2=predicted as infected and cut-off value of 1.5.

### Analysis of prediction accuracy

Sensitivity was calculated to estimate of the model’s ability to detect the presence of infection and specificity as the model’s ability to detect the absence of infection. Accuracy was calculated as the overall prediction ability of the model (Table 3). Sensitivity, specificity, accuracy and respective exact Clopper-Pearson confidence intervals were calculated using MedCalc for Windows, version 18.0 (MedCalc Software, Ostend, Belgium). Cohen’s kappa was calculated in STATA/IC Version 13.as a measure of inter-rate agreement between qPCR (reference test) and NIRS. The PLS scores of the predicted independent samples were analyzed using generalized linear mixed-effects model in STATA/IC Version 13.1. The response variable investigated was the PLS score generated from the PLS calibration models. The effects of infection presence and infection intensity (number of parasite genomes) on the PLS prediction value were investigated. Given that the age of a mosquito may affect NIRS spectra and therewith the PLS score, mosquito age was included as a random effect in the model. Regression coefficients for each factor, confidence intervals and p-values were reported. Model selection was done based on the Akaike information criterion (i.e. the lower the AIC value, the better the model).

**Table 3.** Sensitivity, specificity and accuracy as measures of the performance of a binary classification test TP – True positives; TN – True negative; FP – False positive; and FN – False negatives

| Sensitivity | Specificity | Accuracy |
| --- | --- | --- |
| $\frac{TP}{TP + FN}$ | $\frac{TN}{TN + FP}$ | $\frac{TP + TN}{TP + TN + FP + FN}$ |

## DATA AVAILABILITY

All the data necessary to interpret and replicate the finding on this paper have been made publicly available on the data repository Harvard dataverse (https://doi.org/10.7910/DVN/YD34OX). This includes details on the mean number of parasite genomes of each individual sample; NIR spectra of all the specimens (spc files) with specification to whether they had been included in the training dataset or test dataset for occyst or sporozoite calibration; calibration file (cal file) for oocyst and sporozoite prediction; GRAMS IQ training files (tdfx file) for oocyst and sporozoite prediction; as well as the prediction outputs from IQ Predict for each sample in the test datasets (xls file).

## ACKNOWLEDGEMENTS

The authors acknowledge the Swiss National Foundation of Science for the funding provided to MFM through the Marie-Heim Voegtlin fellowship scheme (PMPDP3-164444) and AXA RF fellowship (14-AXA-PDOC-130) and an EMBO LT fellowship (43-2014) for funding to FB. The authors also wish to thank the Elizabeth Peat and Dorothy Armstrong for the production of mosquitoes at the University of Glasgow and Laura Ciuffreda (supported by EU-FP7 MCSA-ITN Lapaso (607350)) for the ring stage synchronized parasite culture. The authors also thank the Initiative to Develop African Leaders Program (IDeAL) for funding Michelle Muthui and Martin Wagah.

## AUTHORS CONTRIBUTIONS

MFM designed the experiment, cultured the parasites, assisted with the SMFAs, scanned the mosquitoes, analysed the data and drafted the manuscript. MK provided the DNA standards, provided training and commented on the final draft of the manuscript. MM optimized the qPCR method and trained MW. MW performed qPCRs. HF provided mentorship to MFM, was involved in the experimental design and commented on the final manuscript draft. FD provided mentorship to MFM, contributed to the experimental design and data analysis. FB and LRC contributed to the experimental design, setup the parasite culture, led the SMFAs, provided training to MFM in asexual and sexual culture of *Plasmodium falciparum* NF54 as well as contributed to the final manuscript. All authors read and commented on drafts of the manuscript and approved the final version.

