## Supplementary information for "Detection of malaria in insectary-reared *Anopheles gambiae* using near-infrared spectroscopy"

Calibrations were generated using PLS regression by performing multiple leave-one-out cross validations (LOOCV) on two training datasets, each respectively for *Plasmodium falciparum* oocyst and *P. falciparum* sporozoite identification in *Anopheles gambiae* (Keel line) mosquitoes. The spectra that were included in the training data sets have been made available on the data repository Harvard dataverse. The same applies to the GRAMS IQ Version 9.3 file (*.tfdx) and the calibration file (*.cal) that was loaded onto IQ Predict (Add-on of GRAMS IQ Version 9.3). The spectra of the test dataset that were used to validate the calibration with independent samples of unknown infection status has also been made available on Harvard dataverse.

The cross validations of the training datasets showed relatively good prediction accuracy after LOOCV (R=0.65 for the oocyst calibration and R=0.69 for sporozoite calibration) (Figure 1 and 5). The number of latent factors for both calibrations were chosen based on the predicted residual sum of squares curve (PRESS) (Figures 2 and 6) and the PLS beta-coefficients graph (Figures 3 and 7). The spectra residuals were also observed to check for heteroscedasticticity (Figure 4 and 8).

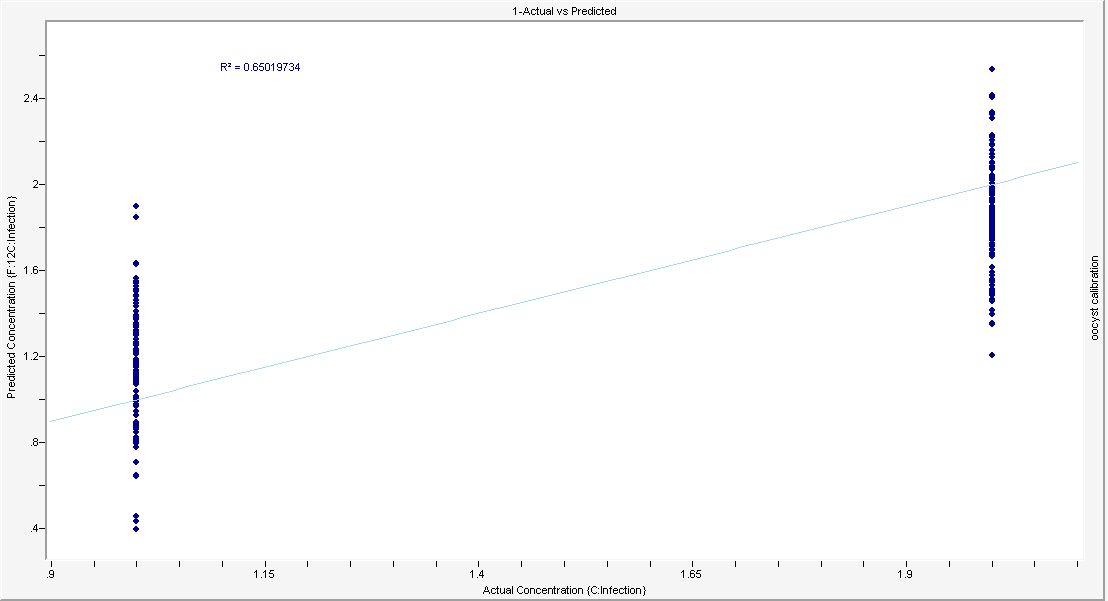

Figure 1. Actual vs Predicted graph. Leave-one-out cross validation to test the self-prediction accuracy of the training dataset used to generate the calibration file to predict the presence of oocyst infection in *Anopheles gambiae* mosquitoes.

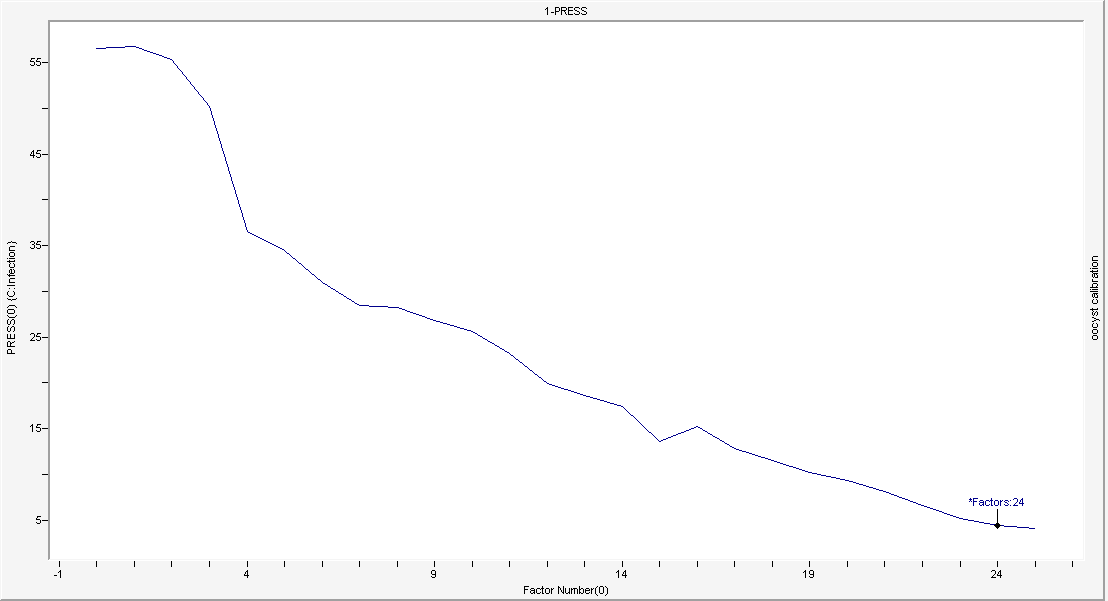

Figure 2 – Prediction residual error sum of squares (PRESS) curve for selection of number of latent factors to be selected for the calibration file for prediction of oocyst infection in *Anopheles gambiae* mosquitoes.

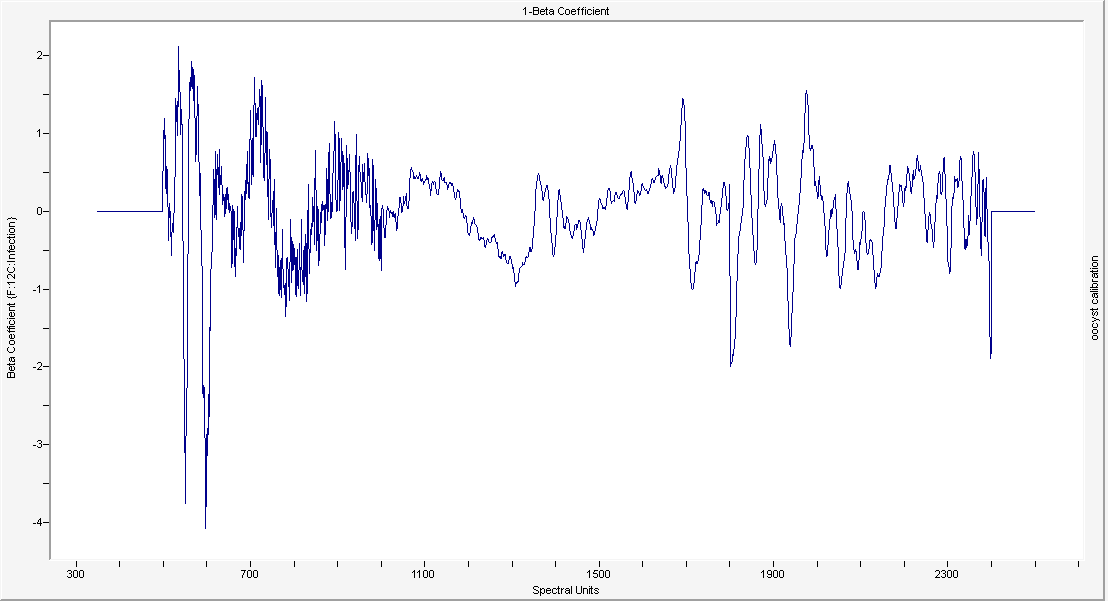

Figure 3– PLS regression beta-coefficients plot used to predict the presence or absence of infection based on 12 latent PLS regression factors. The plots show peaks that determined the differentiation between oocyst-infected and uninfected mosquitoes.

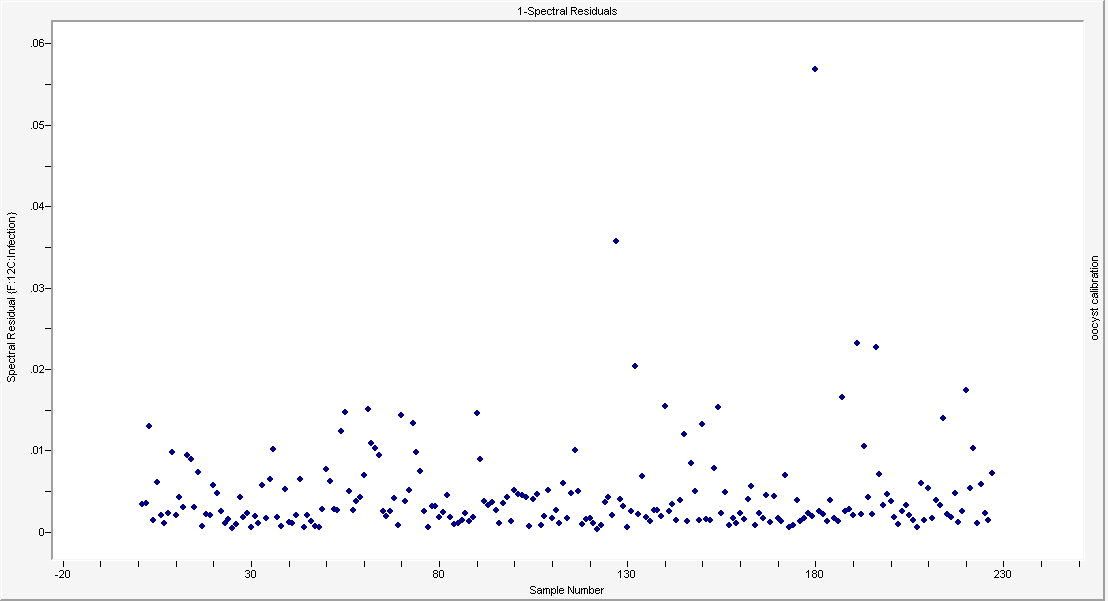

Figure 4 – Spectral residuals of the all the spectra that were self-predicted using LOOCV to form the calibration file to predict oocyst infection in *Anopheles gambiae* mosquitoes.

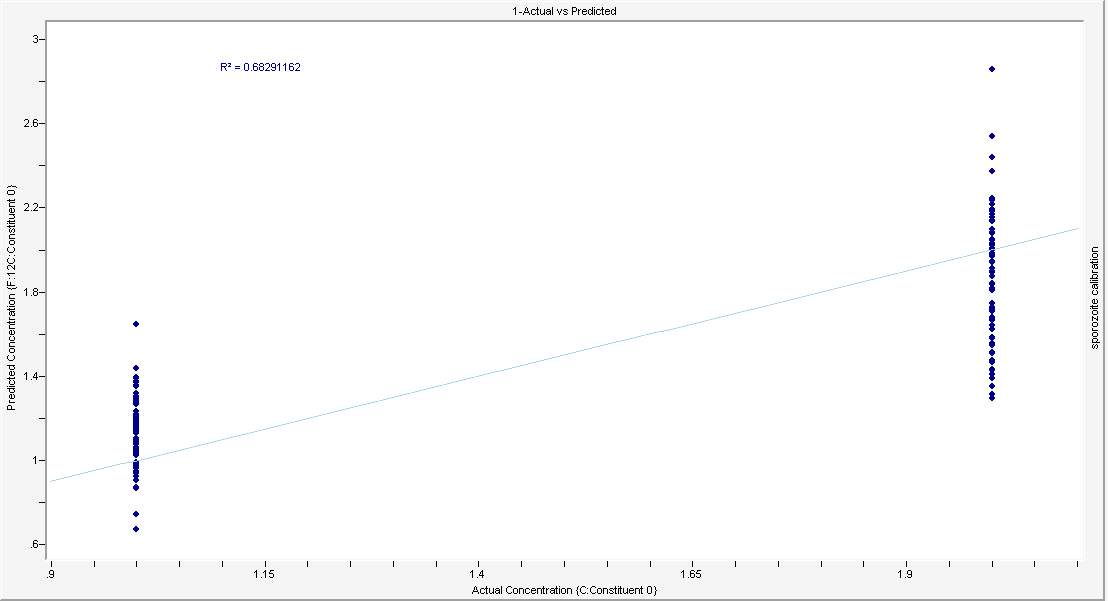

Figure 5. Actual vs Predicted graph. Leave-one-out cross validation to test the self-prediction accuracy of the training dataset used to generate the calibration file to predict sporozoite infection in *Anopheles gambiae* mosquitoes.

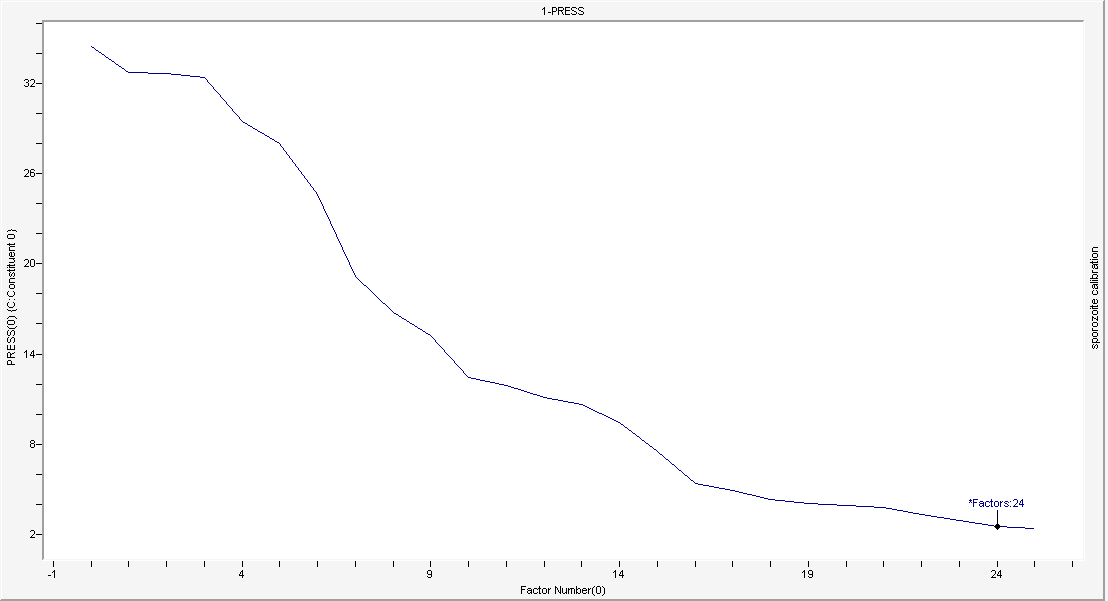

Figure 6 – Prediction residual error sum of squares (PRESS) curve for selection of number of latent factors to be selected for the calibration file for prediction of sporozoite infection in *Anopheles gambiae* mosquitoes.

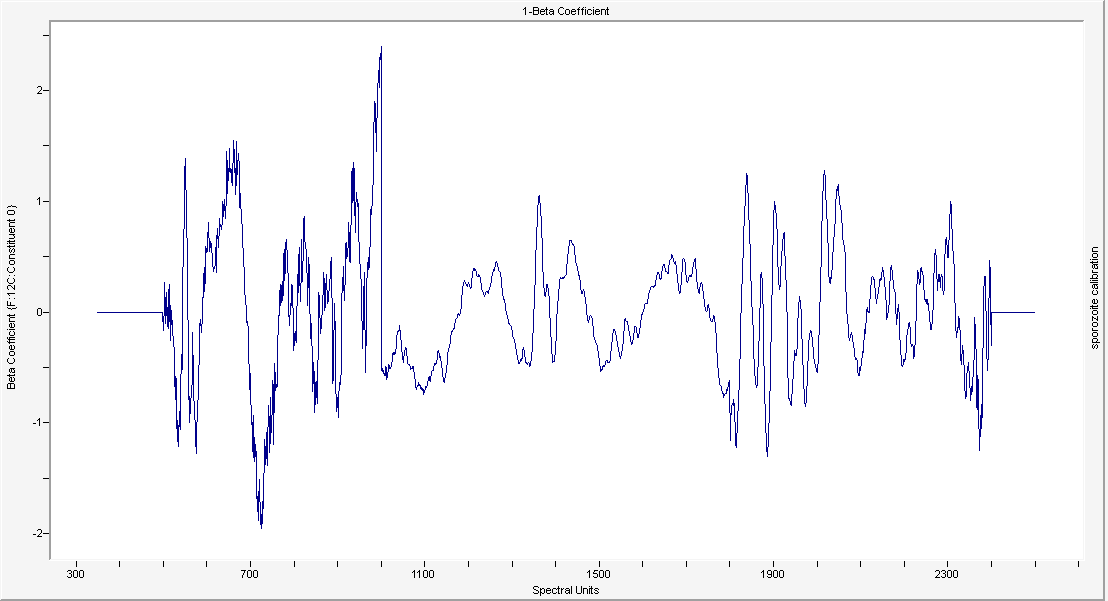

Figure 7 – PLS regression beta-coefficients plot used to predict the presence or absence of infection based on 12 latent PLS regression factors. The plots show peaks that determined the differentiation between oocyst-infected and uninfected mosquitoes.

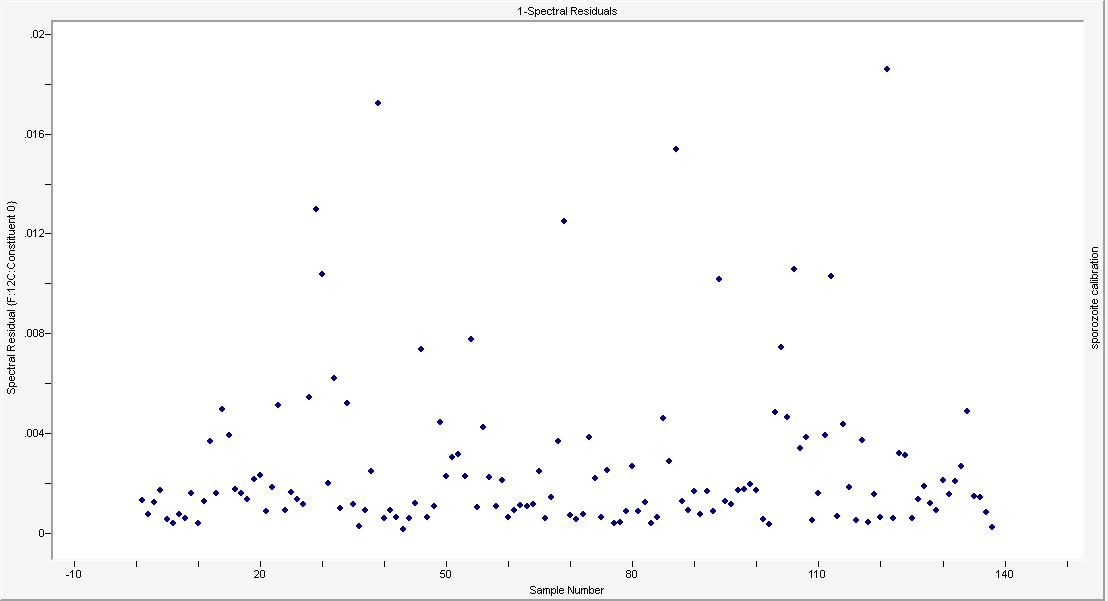

Figure 8 – Spectral residuals of the all the spectra that were self-predicted using LOOCV to form the calibration file to predict sporozoite infection in *Anopheles gambiae* mosquitoes.

The training and test datasets were composed of mosquitoes of different ages and different infection load (Table 1 and Table 2). The mosquitoes were separated by age on the day of the SMFA and kept for either 7 or 14 days to allow oocyst or sporozoite development, respectively.

Table 1 -Number of mosquitoes used in the training and test datasets kept for 7 days to permit oocyst development, by age and parasite load (number of parasite genomes quantified by qPCR. *All uninfected mosquitoes were fed blood containing temperature-inactivated gametocytes in order to reduce the risks of including false PCR-negative mosquitoes in the training and test datasets.

|  | **Mosquito age on the day of SMFA** | | | |
| --- | --- | --- | --- | --- |
|  | **3 days-old** | **4 days-old** | **5 days-old** | **6 days-old** |
| **Training dataset** |  |  |  |  |
| Uninfected* | 7 | 36 | 38 | 25 |
| Less than 200 parasites | 6 | 5 | 5 | 3 |
| 200 to 999 parasites | 16 | 9 | 8 | 5 |
| 1’000 to 9’999 parasites | 18 | 23 | 6 | 5 |
| More than 10’000 parasites | 5 | 5 | 1 | 1 |
| **Test dataset** |  | | | |
| Uninfected* | 4 | 18 | 19 | 12 |
| Less than 200 parasites | 5 | 1 | 4 | 3 |
| 200 to 999 parasites | 3 | 4 | 3 | 0 |
| 1’000 to 9’999 parasites | 8 | 11 | 1 | 2 |
| More than 10’000 parasites | 5 | 3 | 0 | 0 |

Table 2 -Number of mosquitoes used in the training and test datasets kept for 14 days to permit sporozoite development, by age and parasite load (number of parasite genomes quantified by qPCR. *All uninfected mosquitoes were fed blood containing temperature-inactivated gametocytes in order to reduce the risks of including false PCR-negative mosquitoes in the training and test datasets.

|  | **Mosquito age on the day of SMFA** | | | |
| --- | --- | --- | --- | --- |
|  | **3 days-old** | **4 days-old** | **5 days-old** | **6 days-old** |
| **Training dataset** |  |  |  |  |
| Uninfected* | 0 | 23 | 23 | 23 |
| Less than 200 parasites | 2 | 3 | 0 | 0 |
| 200 to 999 parasites | 4 | 2 | 3 | 2 |
| 1’000 to 9’999 parasites | 13 | 6 | 5 | 1 |
| More than 10’000 parasites | 12 | 13 | 3 | 0 |
| **Test dataset** |  | | | |
| Uninfected* | 0 | 7 | 10 | 5 |
| Less than 200 parasites | 0 | 4 | 0 | 0 |
| 200 to 999 parasites | 5 | 0 | 1 | 1 |
| 1’000 to 9’999 parasites | 16 | 3 | 6 | 3 |
| More than 10’000 parasites | 11 | 18 | 1 | 0 |
